# Anti-diuretic activity of a CAPA neuropeptide can compromise *Drosophila* chill tolerance

**DOI:** 10.1101/336057

**Authors:** Heath A. MacMillan, Basma Nazal, Sahr Wali, Gil Y. Yerushalmi, Lidiya Misyura, Andrew Donini, Jean-Paul Paluzzi

## Abstract

For insects, chilling injuries that occur in the absence of freezing are often related to a systemic loss of ion and water balance that leads to extracellular hyperkalemia, cell depolarization, and the triggering of apoptotic signalling cascades. The ability of insect ionoregulatory organs (e.g. the Malpighian tubules) to maintain ion balance in the cold has been linked to improved chill tolerance, and many neuroendocrine factors are known to influence ion transport rates of these organs. Injection of micromolar doses of CAPA (an insect neuropeptide) have been previously demonstrated to improve *Drosophila* cold tolerance, but the mechanisms through which it impacts chill tolerance are unclear, and low doses of CAPA have been demonstrated to cause anti-diuresis in other insects, including dipterans. Here, we provide evidence that low (fM) and high (µM) doses of CAPA impair and improve chill tolerance, respectively, *via* two different effects on Malpighian tubule ion and water transport. While low doses of CAPA are anti-diuretic, reduce tubule K^+^ clearance rates and reduce chill tolerance, high doses facilitate K^+^ clearance from the haemolymph and increase chill tolerance. By quantifying CAPA peptide levels in the central nervous system, we estimated the maximum achievable hormonal titres of CAPA, and found evidence to suggest that CAPA may function as an anti-diuretic peptide in *Drosophila*. We provide the first evidence of a neuropeptide that can negatively affect cold tolerance in an insect, and the first evidence of CAPA as an anti-diuretic peptide in this ubiquitous insect model.

**Summary Statement:** Many insects ion balance in the cold. We show how one neuropeptide can slow ion transport and reduce the cold tolerance of a fly.

## Introduction

The majority of insects are chill susceptible, meaning that they are injured and killed by exposure to temperatures that slow physiological processes without causing ice formation (Baust and Rojas, 1985; MacMillan and Sinclair, 2011a; Overgaard and MacMillan, 2017). There is a growing interest in understanding the biochemical and physiological mechanisms underlying chill susceptibility in ectothermic animals, and several studies have demonstrated that the ability of terrestrial insects to maintain ion and water homeostasis in the cold is closely associated with their chill tolerance (Des Marteaux and Sinclair, 2016; Findsen et al., 2013; Koštál et al., 2006; MacMillan and Sinclair, 2011b; MacMillan et al., 2015d).

In particular, the ability to maintain low extracellular [K^+^] appears critical to chill tolerance (Overgaard and MacMillan, 2017). Chilling slows the activity of membrane bound ATPases (such as Na^+^/K^+^-ATPase), leading to rapid membrane depolarization through a reduction in the electrogenic contribution of these primary active transporters to membrane potential (Andersen et al., 2017a; Djamgoz, 1987; MacMillan et al., 2014; Rheuben, 1972). Over minutes to hours, suppressed ion transport allows for the net movement of Na^+^ and water down their concentration gradients from the haemolymph to the gut. The resulting loss of haemolymph volume, combined with concurrent leak of K^+^ down its concentration gradient from tissues to the haemolymph, can cause progressive haemolymph hyperkalemia (Andersen et al., 2017b; Koštál et al., 2006; MacMillan and Sinclair, 2011b; Overgaard and MacMillan, 2017). As the K^+^ gradient is a critical determinant of cell membrane potentials, this loss of K^+^ balance leads to further cell depolarization, and the combined depolarizing effects of cold and hyperkalemia lead to cell death, likely through triggering of apoptotic signalling cascades (MacMillan et al., 2015c; Yi et al., 2007).

Although chilling can disrupt ion and water balance, leading to organismal injury and death, there is wide variation in chill tolerance among and within insect species, and flies of the genus *Drosophila* are a common and useful model for understanding the mechanisms underlying chill tolerance adaptation and phenotypic plasticity. For example, *Drosophila* species can widely vary in cold tolerance when reared under common-garden conditions; those species that come from more poleward environments are more chill tolerant (Kellermann et al., 2012; MacMillan et al., 2015a) and better maintain K^+^ balance in the cold (Andersen et al., 2017c; MacMillan et al., 2015c). Similarly, *Drosophila melanogaster* acclimated to mild low temperatures (10-15°C) during larval development or as adults are more tolerant of extreme chilling (at 0°C) and better maintain low haemolymph K^+^ during cold stress (Colinet and Hoffmann, 2012; MacMillan et al., 2015b).

Given the above evidence of a role for ion homeostasis in insect chill susceptibility and chill tolerance, there has been recent interest in how the organs responsible for the maintenance of osmotic balance may drive cold tolerance adaptation (Andersen et al., 2017c; Des Marteaux et al., 2018; MacMillan et al., 2015d; Terhzaz et al., 2015; Yerushalmi et al., 2018). Conveniently, the physiology of osmotic balance in *D. melanogaster* is rather well described. The Malpighian tubules of insects (including *Drosophila*) are a single cell thick tubular epithelium, where active transport of ions by V-type H^+^-ATPase and Na^+^/K^+^-ATPase drives the concomitant movement of ions (primarily Na^+^/K^+^ and Cl^-^) and water into the lumen of the tubule to produce an isosmotic primary urine (Dow et al., 1998; O’Donnell, 2008). This primary urine flows into the gut lumen at the midgut-hindgut junction, and active transport in the ileum and rectum allows for the reabsorption of ions and water into the haemocoel and production of a hyperosmotic excreta (Hanrahan and Phillips, 1983; Phillips et al., 1986). The Malpighian tubules of cold adapted and acclimated *Drosophila* better defend rates of fluid and K^+^ secretion at low temperatures, a modification that would help prevent against hyperkalemia (Andersen et al., 2017c; MacMillan et al., 2015d; Yerushalmi et al., 2018). Chill tolerant drosophilids also reduce rectal K^+^ reabsorption during cold stress (preventing hyperkalemia) while those that are chill susceptible have higher rates of K^+^ reabsorption in the cold (which would contribute to hyperkalemia) (Andersen et al., 2017c). Similarly, cold acclimated *D. melanogaster* have reduced rates of K^+^ reabsorption at the rectum relative to warm acclimated flies (Yerushalmi et al., 2018). Although the impacts of cold acclimation on the transport of other ions remain to be examined in *Drosophila*, a recent and thorough analysis with locusts (*L. migratoria*) demonstrated that cold acclimation led to increased rates of rectal Na^+^, Cl^-^and water reabsorption without increasing K^+^ reabsorption (Gerber and Overgaard, 2018). Critically, all of these changes suggest that insect cold tolerance is intimately tied to the ability of the Malpighian tubules and rectum to maintain function at low temperature in a manner that favours low extracellular [K^+^]. All of the above studies, however, have been conducted without the influence of neuroendocrine factors that are known to precisely regulate insect renal function *in vivo*.

Ion and water balance are under tight neuroendocrine control in insects; rates of transport in the Malpighian tubules and hindgut are independently controlled by a variety of factors. Many neuropeptides, for example, are produced in neurosecretory cells in the central nervous system and released into the haemolymph, where they bind to receptors, initiating signaling cascades that alter rates of ion and water secretion or absorption (Coast, 2007). Diuretic factors, such as the corticotropin-releasing factor-related peptide, stimulate fluid secretion by the Malpighian tubules, while anti-diuretic factors can slow rates of primary urine production by the tubules or lead to enhanced reabsorption across the hindgut (Coast et al., 2002). Several factors have been demonstrated to induce diuresis in insects, while the number of described anti-diuretic factors is much more limited, despite widespread appreciation that the ability to slow rates of diuresis is likely critical to insect survival under a wide variety of abiotic conditions (Paluzzi, 2012).

The first member of the CAPA peptide family was originally identified and found to be cardioacceleratory in *Manduca sexta* (Huesmann et al., 1995), and genes encoding these peptides were later discovered in *D. melanogaster*, and called *capability* (Davies et al., 1995; Kean et al., 2002). CAPA peptides have been suggested to have either diuretic or anti-diuretic effects on tubule function, depending on the species under study (Davies et al., 2013; Ionescu and Donini, 2012; Paluzzi, 2012; Pollock et al., 2004). In *Drosophila melanogaster* in particular, CAPA peptides are generally thought to be diuretic, and act through increased nitric oxide, cGMP, and calcium levels in the principal cells of the tubules (Davies et al., 1995; Davies et al., 2013; Kean et al., 2002). By contrast, there is evidence that in the mosquito *Aedes aegypti* (another dipteran), CAPA peptides are instead anti-diuretic, act through cGMP, and counteract the actions of diuretic hormones such as 5-HT and mosquito natriuretic peptide (i.e. a DH_31_–related peptide) (Ionescu and Donini, 2012; Sajadi et al., in press). Critically, these anti-diuretic effects observed in *A. aegypti* occur at very low (e.g. femtomolar) concentrations of CAPA peptides, while higher supraphysiological doses (e.g. 100 µM) were instead found to have modest diuretic effects (Ionescu and Donini, 2012). Considering these recent findings from a relatively closely-related insect led us to reconsider the question of whether low concentrations of CAPA peptide are indeed capable of causing anti-diuresis in *D. melanogaster*, as it does in other insects (Coast et al., 2010; Paluzzi et al., 2008; Sajadi et al., in press).

Whether CAPA peptides are diuretic or anti-diuretic is of great importance to understanding neuropeptide control of salt balance alone, but it is also critical background knowledge if we are to understand the mechanisms through which abiotic stressors, such as desiccation or cold stress, impact organismal fitness. Given their role in osmotic balance, and the importance of osmotic balance to both desiccation and chill tolerance, members of the CAPA peptide family have already been tested in such a context (Terhzaz et al., 2015). In *Drosophila*, cold stress leads to upregulation of *capa* mRNA; injections (µM) of *Manse*-CAP2b speed up recovery from chill coma, and targeted knockdown of the *capability* gene slows chill coma recovery (Terhzaz et al., 2015). The physiological means by which CAPA has these effects on chill tolerance, however, remain unknown.

Here, we test the hypothesis that CAPA peptide exerts differential activity on chill tolerance in *D. melanogaster* through dose-dependent effects of this neuropeptide on Malpighian tubule ion and water secretion. We predicted that very low (e.g. fM) doses of CAPA would be anti-diuretic (inhibit fluid secretion by the Malpighian tubules), which would impair K^+^ clearance from the haemolymph and thereby impair chill tolerance. We further predicted that higher doses (µM) of CAPA would be diuretic (as previously demonstrated), which would improve chill tolerance through increased K^+^ clearance by the tubules. We follow this mechanistic analysis by addressing a simple question: What levels of CAPA peptides are likely to occur in the haemolymph of a free-living adult fly?

## Materials and Methods

### Animal husbandry

The population of *Drosophila melanogaster* used in this study was derived from isofemale lines collected in southwestern Ontario, Canada in 2007 (Marshall and Sinclair, 2010). All flies were reared at 25°C (14 h:10 h light:dark cycle) in 200 mL bottles containing 50 mL of Bloomington *Drosophila* medium (Lakovaara, 1969). Groups of ∼80 adult flies were given access to fresh food for 2-3 h to oviposit before being removed to ensure rearing densities of ∼100 eggs per bottle. Females were collected under brief CO_2_ anaesthesia (<2 min exposure to CO_2_) upon final ecdysis and transferred to vials containing 7 mL of the same medium at a density of 20 flies per vial, where they were left to mature for seven days, in part to avoid effects of anaesthesia on chill tolerance (Nilson et al., 2006). All experiments were thus conducted on seven day-old virgin female flies.

### Short-term effects of CAPA on chill coma recovery

To examine the effects of CAPA injection on chill coma recovery time (CCRT), we conducted a dose-response experiment. Individual female flies (n=9-14 flies per treatment group) were transferred (without anaesthesia) to 4 mL glass screw top vials that were submerged in an ice-water slurry (0°C), which rapidly induced chill coma as it is below the critical thermal minimum temperature for this population of *D. melanogaster* (MacMillan et al., 2017). Flies were left at 0°C for either 30 min or 2 h, whereupon they were individually removed from the ice-water and placed on a plasticine surface on top of a double-walled glass plate held at 0°C. A mixture of ethylene glycol and water was circulated through the glass dish from a refrigerated circulating bath (MX7LL, VWR International, Mississauga, Canada) to keep the flies in chill coma during injection.

Solutions containing an insect CAPA neuropeptide (*Aedes aegypti* CAPA-2, *Aedae*CAPA-2: pQGLVPFPRV-NH_2_), were prepared in order to achieve a final post-injection concentration of 10^−15^, 10^−12^, 10^−9^, and 10^−6^ M based on a ∼80 nL haemolymph volume (Folk et al., 2001). CAPA peptide solutions were made up in a 1:1 mixture of *Drosophila* saline (117 mM NaCl, 20 mM KCl, 8.5 mM MgCl_2_, 2 mM CaCl_2_, 10 mM glutamine, 20 mM glucose, 4.3 mM NaH_2_PO_4_, NaHCO_3_, 15 mM MOPS, pH 7.0) and Schneider’s insect medium (Sigma-Aldrich, Oakville, ON, Canada). Injections (18.4 nL) were administered into the haemolymph at the base of the left wing (mesopleurum) using a pulled glass microcapillary connected to a Nanoject system (Drummond Scientific Co., Broomall, USA). Control flies were put on the cooled plate but received no injection, whereas sham-injected flies were injected with saline that did not contain any CAPA peptide. Flies were subsequently placed back into their vials and re-submerged in the ice-water at 0°C. All flies spent a total of 3 h at 0°C before being removed to record chill coma recovery time, and thus the groups injected at 30 min and 2 h differed only in the time during the cold stress at which the injection was given.

Chill coma recovery time (CCRT) was measured as previously described (MacMillan et al., 2015b). Briefly, flies, in their glass vials, were removed from the cold and placed on a laboratory bench lined with paper at 23°C. The flies were observed in their vials without being disturbed, and the time taken for each fly to right itself and stand on all six legs was recorded.

### Effects of CAPA peptide injection on chill coma recovery after prolonged chilling

To determine whether CAPA injection similarly impacted chill coma recovery after chronic chilling, we measured chill coma recovery following 16 h exposure to 0°C (n=18-19 flies per treatment group). As in the previous experiment, *D. melanogaster* females were individually separated into 4 mL glass vials that were submerged in an ice-water slurry (0°C). After 15 hours at 0°C, flies were injected with 18.4 nL of either the sham, or CAPA peptide (to achieve final concentration of 10^−6^ M or 10^−15^ M, the lowest and highest doses used in the prior short term chill experiment). Control animals were handled in the same way as described above but received no injection. The flies were then returned to their vials and placed in the ice water slurry for a further 1 h (meaning they experienced 16 h at 0°C in total), at which point the flies were removed from the ice bath and transferred to room temperature to measure CCRT as above.

### Effects of CAPA peptide injection on survival following a cold stress

To examine whether CAPA peptide injection influences survival following prolonged chilling, we injected flies early in a 16 h cold stress and recorded survival outcomes (n=50 flies per treatment group). As above, individual flies were placed into 4 mL glass vials and submerged in an ice-water mixture (0°C). Flies were injected after 1 h at 0°C with either the saline alone (i.e. sham), or saline containing CAPA peptide in order to achieve a final titre of 10^−6^ M or 10^−15^ M in the haemolymph. The flies were then returned to their vials, and held at 0°C for a further 15 h (16 h at 0°C in total) upon which they were removed from the cold and transferred to 40 mL plastic vials containing 7 mL of fresh food medium in groups of 10. The flies were then held under their rearing conditions (25°C) for 24 h, before being visually inspected to determine survival, which was scored for each fly on a 5-point scale: 1 = dead (no movement), 2 = moving but unable to stand, 3= standing but not climbing, 4 = climbing, 5 = flying.

### Effects of CAPA and cGMP on Malpighian tubule function

We measured the effects of low (10^−15^ M) and high (10^−6^ M) doses of CAPA as well as low (10^−8^ M) and high (10^−3^ M) cGMP on Malpighian tubule fluid and ion secretion rates using Ramsay assays combined with the ion-selective electrode technique, as previously described (MacMillan et al., 2015d). Flies (CAPA: n=25-32 per treatment group; cGMP: n=12-13 per treatment group) were dissected under *Drosophila* saline to carefully isolate the anterior pair of Malpighian tubules, which were separated from the gut at the ureter. The pair of tubules connected at the ureter were transferred to a Sylgard-lined petri dish with ∼4 mm deep wells that were set 0.5 cm apart. The dish was filled with hydrated paraffin oil to prevent sample evaporation. A 20 µL droplet of a 1:1 mixture of Schneider’s insect medium and *Drosophila* saline was added to each well. This mixture either contained no neuropeptide or cGMP (i.e. control), contained 10^−6^ M or 10^−15^ M CAPA, or 10^−8^ M or 10^−3^ M 8-bromo cGMP (Sigma-Aldrich, Oakville, ON, Canada), a membrane-permeable analog of cGMP with greater resistance to phosphodiesterases compared to its parent compound. One pair of tubules was placed in the drop of bathing medium and the proximal end of one tubule was pulled out of the drop and wrapped around a minuten pin. As the tubule remaining in the bathing solution actively secretes fluid, a droplet forms at the ureter. After 30 min, the droplet was detached from the ureter, lifted off of the Sylgard surface and its diameter was measured with an eyepiece micrometer. Droplet volume was calculated from the diameter of the secreted droplet

Concentrations of ions in the primary urine secreted by the Malpighian tubules were measured using the ion-selective microelectrode technique (Rheault and O’Donnell, 2004). Ion-selective microelectrodes were pulled from glass capillaries (TW150-4; World Precision Instruments, Sarasota, FL, USA) using a P-97 Flaming Brown micropipette puller (Sutter Instruments Co., San Rafael, CA, USA) to produce a probe with a short shank and wide angle with a ∼5 µm tip diameter. Micropipettes were then silanized at 300°C with N,N-dimethyltrimethylsilylamine (Fluka, Buchs, Switzerland). For K^+^ measurements, the micropipette was back filled with 100 mM KCl and front filled with K^+^ ionophore (potassium ionophore I cocktail B; Fluka, Bachs, Switzerland). For Na^+^ measurements, the micropipette was backfilled with 100 mM NaCl and front filled with Na^+^ ionophore (sodium ionophore II cocktail A; Fluka, Buchs, Switzerland). Ion-selective microelectrodes were dipped in a solution of polyvinylchloride in tetrahydrofuran to prevent the ionophore from leaking out of the microelectrode upon contact with the paraffin oil. The circuit was completed with a reference electrode, pulled from glass capillaries (IB200F-4, World Precision Instruments, Sarasota, FL, USA), and backfilled with 500 mM KCl. Both electrodes were connected to an amplifier (ML 165 pH Amp), which was connected to PowerLab 4/30 data acquisition system (AD Instruments Inc., Colorado Springs, USA). Data were recorded using Labchart 6 Pro software (AD Instruments Inc.).

Ion concentrations (mM) in the secreted droplets were calculated using the following equation (Donini *et al.,* 2008):

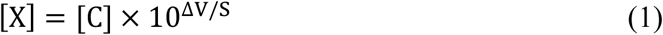

where [X] is the concentration of the secreted fluid droplet, [C] is the ion concentration of one of the standards, ΔV is the voltage difference between the secreted fluid droplet and the voltage measured in the same standard, S is the difference in voltage between two standard solutions (which cover a 10-fold difference in ion concentration). Rates of ion secretion from the tubule were then calculated from droplet concentrations and fluid secretion rates.

### Quantification of CAPA peptides in the *D. melanogaster* haemolymph and nervous system

Haemolymph was extracted from flies (n=60 flies per sample) as previously described (MacMillan and Hughson, 2014), pooled into methanol:acetic acid:water (90:9:1) and frozen for later processing. The thoracicoabdominal ganglion was dissected from adult female *D. melanogaster* using forceps under *Drosophila* saline in the view of a dissecting microscope. Each dissection took approximately two minutes and ganglia were transferred to a microcentrifuge tube containing methanol:acetic acid:water (90:9:1) held on ice as they were individually dissected to produce three biological replicates, each containing 20 ganglia, which were stored at −80°C for later analysis. Peptidergic extracts were then isolated by sonicating ganglionic samples on ice for two consecutive 5 s pulses using an XL 2000 Ultrasonic Processor (Qsonica LL, Newtown, CT, USA). Ganglionic and haemolymph homogenates were then centrifuged at 10,000 × *g* for 10 min at 4°C. The supernatants were transferred to a new microcentrifuge tubes, dried in a Jouan RC10 series vacuum concentrator (Jouan Inc., Winchester, VA, USA) and reconstituted in 0.4% trifluoroacetic acid (TFA). Samples were then applied to C18 Sep-Pak cartridges (Waters Associates, Mississauga, ON, Canada) following sequential washing and equilibration with 10 mL of acetonitrile (ACN), 5 mL of 50% ACN, 0.5% acetic acid (HAcO), and finally 5 mL 0.1% TFA. To ensure complete binding of peptidergic extracts, samples were passed through the Sep-Pak cartridge at least three times. Once the samples were loaded, the cartridges were first washed/desalted with 5 mL 0.1% TFA and subsequently with 5 mL 0.5% HAcO to remove TFA. Samples were then eluted with 2 mL each of 0.5% HAcO, 10%, 20%, 30%, 40% and 50% ACN all containing 0.5% HAcO. The eluants were dried in a vacuum concentrator as above and samples were then stored at −20°C for later quantification analysis using an enzyme-linked immunosorbent assay (ELISA).

A CAPA peptide-specific ELISA was developed based on an earlier report describing a crustacean cardioactive peptide (CCAP)-specific ELISA used in the stick insect, *Baculum extradentatum* (Lange and Patel, 2005). A rabbit anti-CAPA affinity purified polyclonal antibody (a generous gift from Prof. Ian Orchard, University of Toronto Mississauga) was diluted 1:1000 in carbonate buffer (15 mM Na_2_CO_3_, 35 mM NaHCO_3_, pH 9.4) and 100 µL of this antibody solution was applied into each well of a high-binding 96-well ELISA plate (Sarstedt Inc., Montreal, QC) and incubated overnight at 4°C. The following day, wells of the ELISA plate were washed three times with 250 µL wash buffer (346 mM NaCl, 2.7 mM KCl, 1.5 mM KH_2_PO_4_, 5.1 mM NaH_2_PO_4_, 0.5% Tween-20) and after the final wash, each well was loaded with 250 µL block solution (phosphate buffered saline containing 0.5% (w/v) each of skim milk powder and protease-free bovine serum albumin) and incubated for 1.5-2 h at room temperature (RT). During the blocking step, standards and unknowns were prepared as follows: commercially synthesized *D. melanogaster* CAPA2 (*Drome*CAPA-2: ASGLVAFPRV-NH_2_) peptide and an N-terminally biotinylated *Drome*CAPA2 (biotin-CAPA-2) peptide analog (Genscript Inc., Piscataway, NJ) were prepared in block solution. Serial dilutions of synthetic *Drome*CAPA-2 ranging between 2.5 fmol/100 µL and 50 pmol/100 µL were loaded in triplicate onto the ELISA plate along with block solution alone applied to wells with and without antibody coating (for maximum signal and blank/background controls, respectively). Once standards were loaded, aliquots of the dried ganglionic fractions or haemolymph samples from above resuspended in block solution were loaded (100 µL/well) in triplicate onto the ELISA plate. The standards and unknowns were incubated at RT for 1.5 h and then 100 fmol of biotin-*Drome*CAPA-2 was dispensed to each well already containing the standards or unknown samples and the plate was incubated overnight at 4°C on a bidirectional rocking platform. The next day, contents in the wells were discarded and the plate was washed four times with wash buffer (250 µL/well). After the last wash was discarded, wells were loaded with 100 µL Avidin-HRP conjugate (Bio-Rad, Mississauga, ON) diluted 1:2000 in block buffer and incubated for 1.5 h at RT. Following this incubation, contents in the wells were discarded and the plate was washed three times with wash buffer (250 µL/well). After the final wash solution was discarded, each well was loaded with 3,3′,5,5′-tetramethylbenzidine (TMB) liquid horseradish peroxidase substrate and incubated for approximately 10 min at RT to allow blue end product development. Without discarding the TMB solution, each well then received 100 µL of 2N HCl to stop the reaction and the absorbance was then measured at 450 nm using a Synergy 2 Multi-Mode Microplate Reader (BioTek, Winooski, VT, USA).

### Data analysis

All data analysis was completed in the R environment for statistical computing, version 3.4.3 (R Development Core Team, 2017). Chill coma recovery time following 3 h at 0°C was compared among treatment groups and injection times using a two-way ANOVA. Chill coma recovery times following prolonged chilling were not normally distributed (Shapiro-Wilk test; W=0.94, P=0.002), so a Kruskal-Wallis test was used to test for effects of injection treatment (10^−15^ M and 10^−6^ M CAPA) on chill coma recovery following 16 h at 0°C. Similarly, a Kruskal-Wallis test was used to test for effects of injection treatment on locomotor function and survival scores among flies following 16 h at 0°C. In both cases, pairwise comparisons were completed with a Wilcoxon rank sum test. Effects of CAPA peptide or cGMP concentration on Malpighian tubule fluid, Na^+^ and K^+^ secretion rates, and [Na^+^] and [K^+^] in the secreted fluid were tested using ANOVA followed by Tukey’s HSD (if data were normally distributed) or were conducted on Kruskal-Wallis tests, followed by Wilcoxon rank sum tests (if the data were not normally distributed). In all cases, post-hoc analyses were corrected for multiple comparisons (Benjamini and Hochberg, 1995). All values reported in the results are mean ± sem unless otherwise stated.

## Results

### Short-term effects of CAPA on chill coma recovery

The effects of CAPA injection during a short (3 h) chilling stress on chill coma recovery time (CCRT) were strongly dose-dependent (Figure 1), and sham injected flies recovered from chill coma slightly later than those given no injection (Two-way ANOVA: F=123.4, *P*<0.001). Flies injected with the lowest dose of CAPA peptide (10^−15^ M) recovered from chill coma 3.65 min (or 33%) more slowly than sham injected flies, while those injected with the highest dose (10^−6^ M) recovered 1.94 min (or 20%) faster than sham injected flies, on average. Flies injected with CAPA peptide 30 min into the cold stress recovered more quickly than those injected 2 h into the cold stress (F=51.6, *P*<0.001), regardless of the dose applied, but the same trend was observed in sham and even control flies (that received no injection), indicating that this effect of timing is likely an artefact and that the timing of CAPA injection has little effect on CCRT (Figure 1 insert). The dose of CAPA applied and the timing of injection did, however, significantly interact to impact CCRT (two-way ANOVA: F=2.4, *P*=0.042). This modest interaction appears to be driven by a somewhat larger effect of early injection on flies given the highest dose (10^−6^ M) of CAPA (Figure 1).

**Figure 1.**
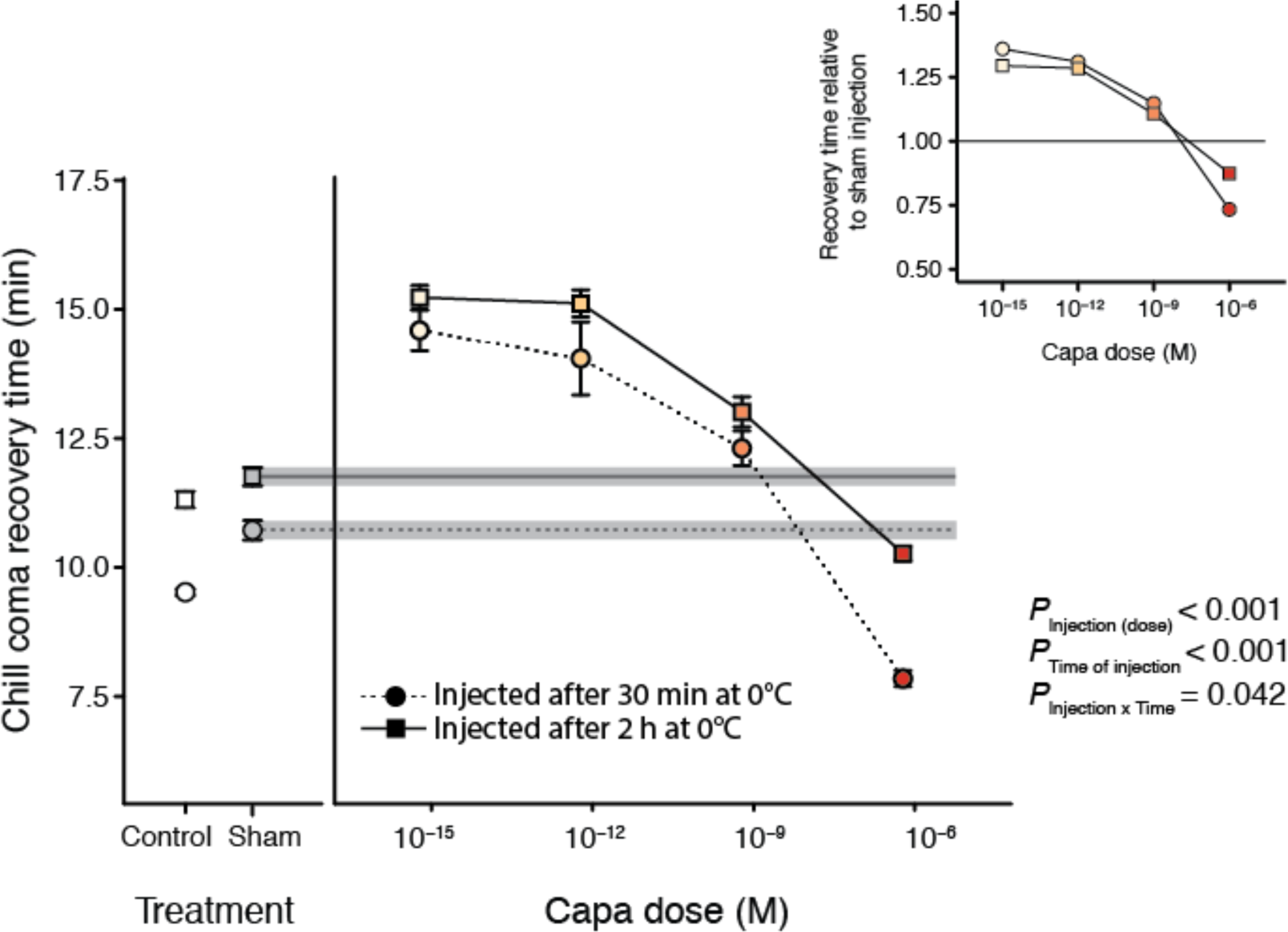
Dose-responsive effects of CAPA injection on chill coma recovery time of adult female *D. melanogaster*. All flies were held individually at 0°C for a total of 3 h, and were injected either 30 min (circles with dashed line) or 2 h (squares with solid line) into the cold stress. Flies were injected with a dose of CAPA peptide in saline to achieve 10^−15^, 10^−12^, 10^−9^, 10^−6^ M CAPA in the haemolymph. Sham flies were given an injection of saline only (without CAPA) whereas control flies were not injected. n=9-14 flies per treatment group.

### Effects of CAPA peptide injection after prolonged chilling on chill coma recovery and survival

CAPA peptide injection also had significant effects on chill coma recovery following prolonged cold exposure (Figure 2A, Kruskal Wallis test: *H* = 22.2, *P*<0.001). Flies given a sham injection of saline before recovering from 16 h at 0°C did not significantly differ in chill coma recovery time from control flies given no injection (*P*=0.447). Flies that were injected with a low dose (10^−15^ M) of CAPA peptide recovered significantly more slowly than control (*P*=0.010) and sham injected flies (*P*=0.021). By contrast, flies injected with a high dose (10^−6^ M) of CAPA recovered more quickly from chill coma than both control flies (*P*=0.047) and sham injected flies (*P*=0.010). A separate set of flies injected in the same manner (15 h into a 16 h exposure to 0°C) were observed 24 h after injection. CAPA injection significantly affected the incidence of chilling injury and death following this prolonged cold exposure (Figure 2B, Kruskal-Wallis test: *H*=22.9, *P*<0.001). Flies injected with a low dose (10^−15^ M) of CAPA peptide had lower survival scores than those given a sham injection (*P*=0.011), while those given a high dose (10^−6^ M) were more likely to be uninjured 24 h after removal from the cold (*P* = 0.019). The median survival score for a fly injected with a low dose (10^−15^ M) of CAPA peptide was 2 (a fly that was moving but unable to stand) while that of a fly injected with a high dose (10^−6^ M) was above 4 (A fly that can climb with coordination and potentially fly).

**Figure 2.**
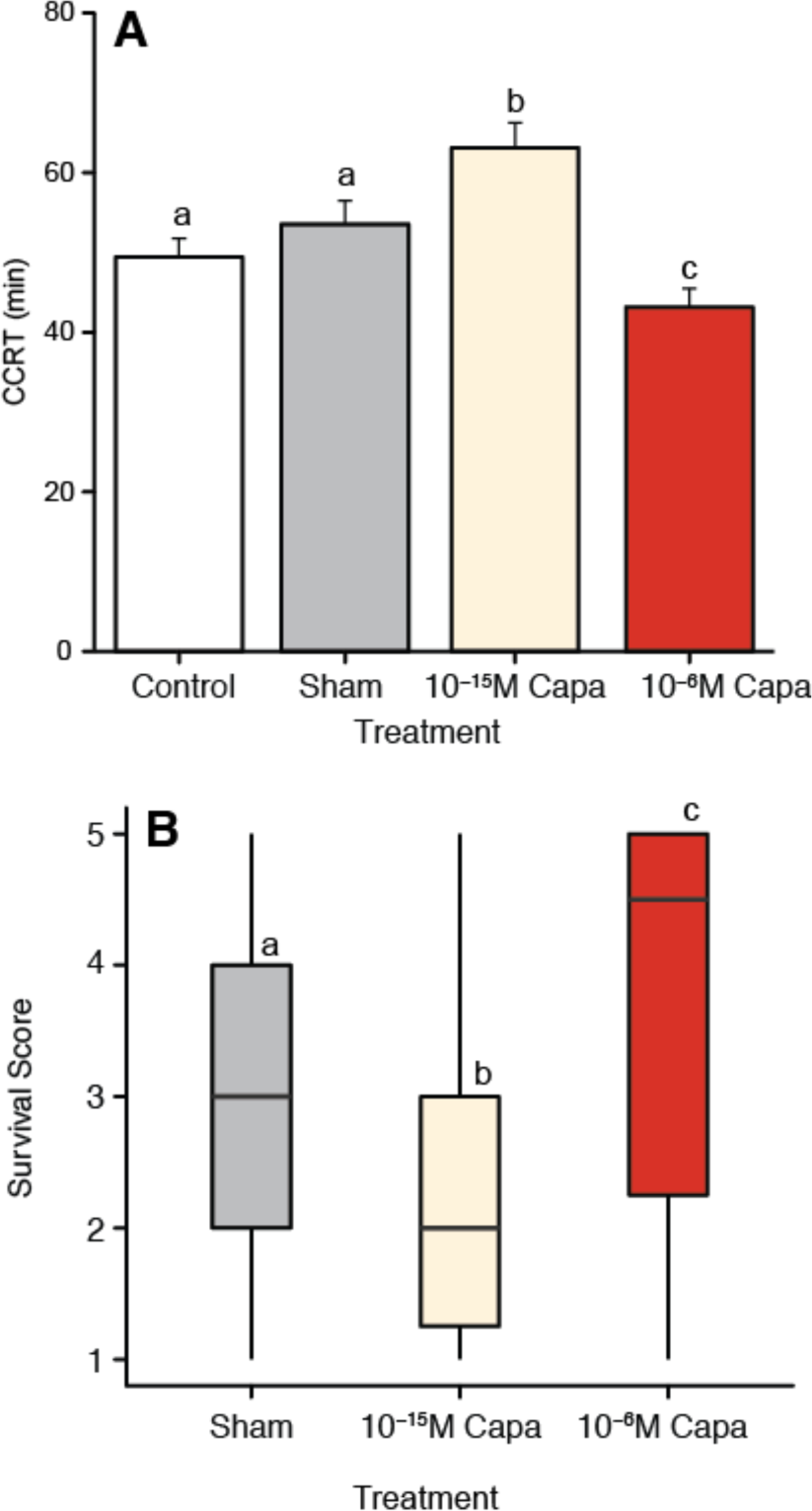
The dose-dependent effects of CAPA peptide injection on chill coma recovery time and survival following prolonged cold stress in adult female *D. melanogaster*. A) Mean ± sem chill coma recovery time of flies exposed to 16 h at 0°C. B). Boxplot of survival scores of flies following 16 h at 0°C. Survival was scored on a five point scale, with a score of 5 being a fly that is able to stand, walk in a coordinated manner, and initiate flight, and 1 being a fly showing no signs of life (see methods for details). The central horizontal line indicates the median value, the box represents the inter-quartile range, and vertical lines denote the full range of the data. In both experiments, flies were injected 15 h into a cold stress of 16 h at 0°C. Flies were injected with saline containing CAPA peptide to achieve an effective circulating titer of 10^−15^ or 10^−6^ M in the haemolymph, were given a sham injection of saline only or received no injection (control). Bars or boxes that share a letter within a panel do not significantly differ.

### Effects of CAPA on Malpighian tubule function

To test whether the observed effects of CAPA peptide on cold tolerance were driven by effects on Malpighian tubule function we directly measured tubule fluid and ion secretion rates using Ramsay assays. The dose of CAPA applied to tubules significantly altered rates of primary urine production (Figure 3A; *H*=8.3, *P*=0.016). Specifically, tubules treated with 10^−15^ M CAPA had approximately 28% lower secretion rates relative to both control (saline only; *P*=0.032) and a higher dose (10^−6^ M) of CAPA peptide (*P*=0.024). The dose of CAPA applied did not affect [Na^+^] of the secreted fluid (Figure 3B; *H*=2.1, *P*=0.348), but strongly affected [K^+^] in the secreted fluid (Figure 3C; *H*=18.0, *P*<0.001). Tubules bathed in 10^−6^ M CAPA peptide produced fluid with significantly higher [K^+^] than both control tubules (42% higher, *P*<0.001) and those bathed in the lower dose of CAPA peptide (34% higher, *P*=0.005), while fluid from tubules bathed in 10^−15^ M CAPA and control tubules did not differ in [K^+^] (*P*=0.849). Using fluid secretion rates and measured ion concentrations, we quantified rates of ion secretion by the tubules during the assay. The dose of CAPA peptide applied had significant effects on the secretion of both Na^+^ (Figure 3D; *H*=8.5, *P*=0.014) and K^+^ ions (Figure 3E; *H*=10.1, *P*=0.006). In the case of Na^+^, tubules bathed in the lower dose (10^−15^ M) of CAPA peptide secreted less Na^+^ than control tubules (*P*=0.021) or those bathed in 10^−6^ M CAPA (*P*=0.043). Tubules bathed in 10^−15^ M CAPA secreted significantly less K^+^ than those bathed in 10^−6^ M CAPA (*P*=0.004), and K^+^ secretion from control tubules was intermediate between the two CAPA treatments (*P*>0.05 in both cases). Using rates of Na^+^ and K^+^ secretion, we calculated the ratio of these two ions secreted from the tubules (Na^+^:K^+^ ratio), and this ratio was significantly impacted by the dose of CAPA peptide administered (Figure 3F; *H*=9.4, *P*=0.009). Malpighian tubules exposed to 10^−6^ M CAPA had a significantly lower Na^+^:K^+^ ratio than control tubules *P*=0.008), and tubules exposed to 10^−15^ M CAPA did not differ from either the control (P=0.306) or the 10^−6^ M CAPA peptide treatment (*P*=0.072).

**Figure 3.**
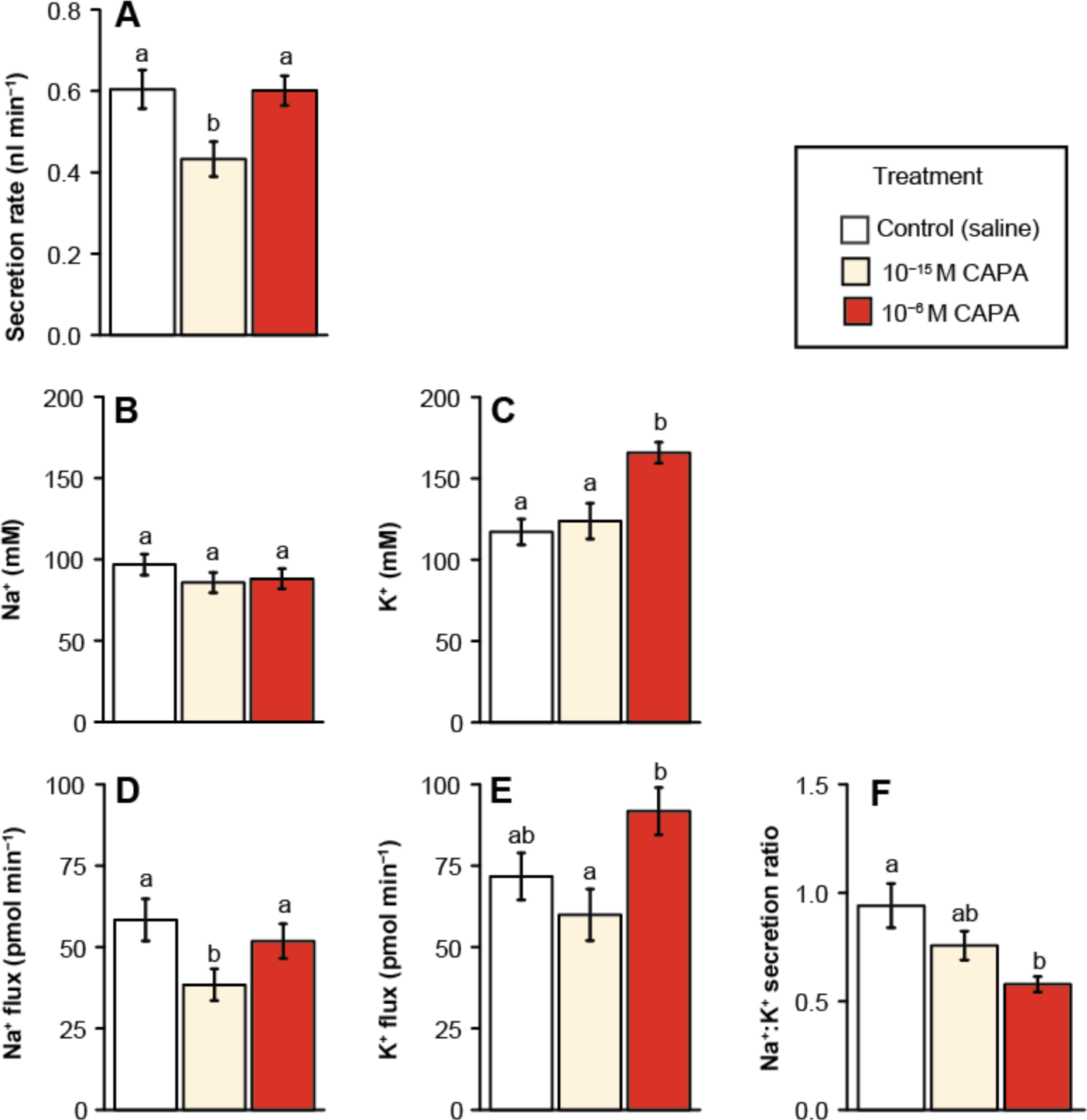
Effects of CAPA peptide on *in vitro* fluid and cation (Na^+^ and K^+^) secretion rates of unstimulated Malpighian tubules of adult female *D. melanogaster*. 1 fM (10^−15^ M) and 1 µM (10^−6^ M) of CAPA peptide were applied to otherwise unstimulated tubules. Data shown are rates of fluid secretion by the tubules (A) as well as concentrations of Na^+^ (B) and K^+^ (C) measured in the secreted droplets, rates of Na^+^ (D) and K^+^ (E) flux expressed independently of water flux, and the ratio of Na^+^:K^+^ (F) in the secreted fluid. In all cases values presented are mean ± sem. Bars that share a letter within a panel do not significantly differ.

### Effects of cGMP on Malpighian tubule function

We tested for effect of two doses of 8-bromo-cGMP on the function of *Drosophila* tubules. The dose of cGMP applied significantly impacted rates of primary urine production (Figure 4A; *F*=3.8, *P*=0.030). Tubules bathed in 10 nM (10^−8^ M) cGMP had reduced rates of secretion relative to control tubules (*P*=0.034), whereas tubules exposed to 1 mM cGMP (10^−3^ M) had similar rates of secretion to control tubules (*P*=0.893). As with CAPA peptide treatments, we found that the dose of cGMP did not impact [Na^+^] (Figure 4B; *F*=0.4, *P*=0.673) or [K^+^] in the secreted fluid (Figure 4C; *F*=2.4, *P*=0.103). The dose of cGMP applied significantly impacted Na^+^ secretion rates (Figure 4D; *F*=4.3, *P*=0.021) and tended (nearly significantly) to impact K^+^ secretion rates (Figure 4E; *F*=3.2, *P*=0.051). Post-hoc analyses revealed that tubules exposed to a low dose of cGMP (10^−8^ M) secreted significantly less Na^+^ (*P*=0.027) and K^+^ (*P*=0.041) than control tubules, while those exposed to the higher dose of cGMP (10^−3^ M) did not differ from the control rates of secretion of either ion (Na^+^: *P*=0.948; K^+^: *P*=0.316). Exposure to cGMP significantly impacted the ratio of Na^+^ and K^+^ secreted by the Malpighian tubules (H=7.6, *P*=0.022), but none of the post-hoc pairwise comparisons revealed significant differences among the groups (Figure 4F; *P*>0.05 in all cases).

**Figure 4.**
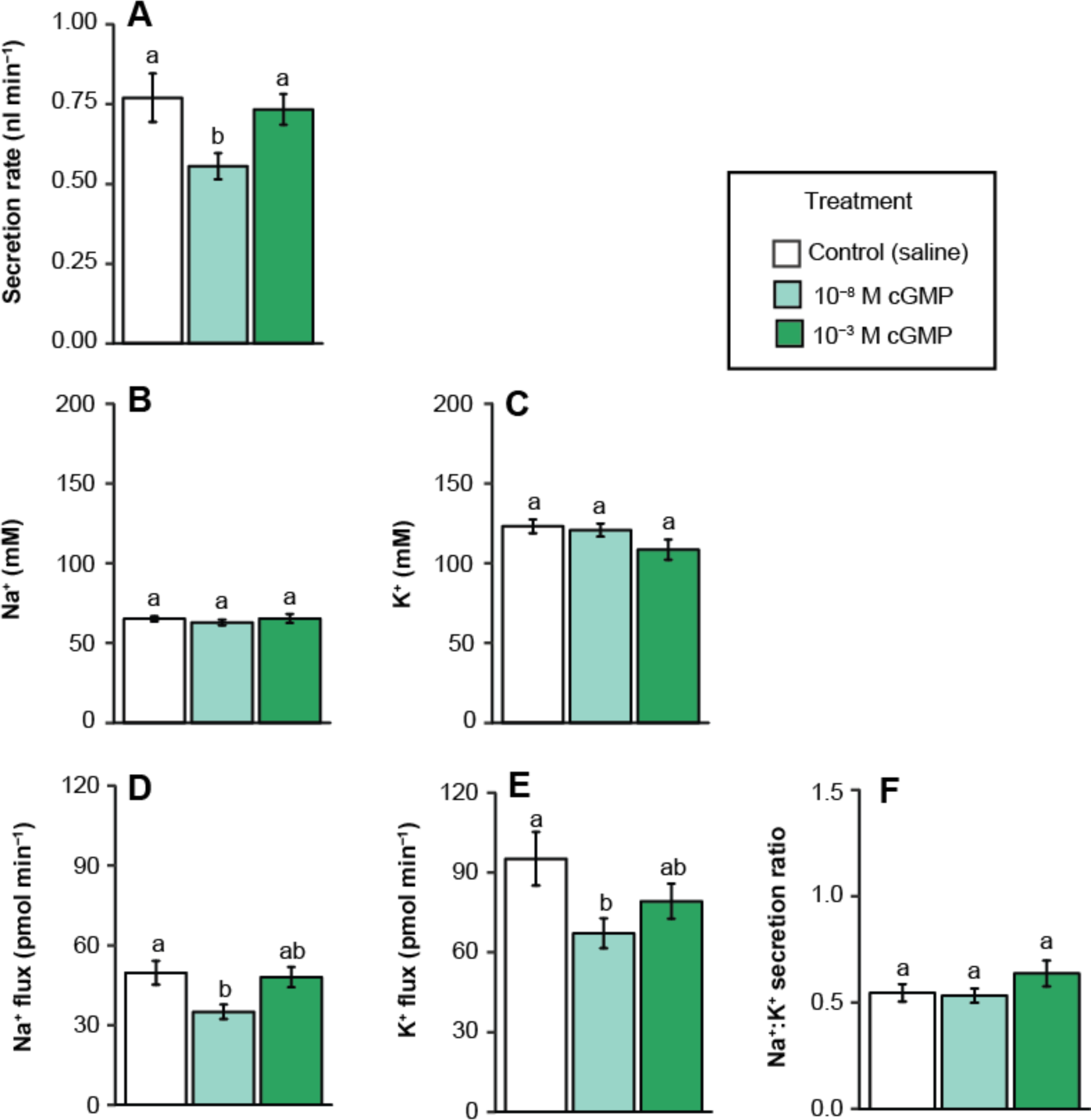
Effects of cGMP on *in vitro* fluid and cation (Na^+^ and K^+^) secretion of the Malpighian tubules of adult female *D. melanogaster*. 10 nM (10^−8^ M) and 1 mM (10^−3^ M) doses of cGMP were applied to otherwise unstimulated tubules. Data shown are rates of fluid secretion by the tubules (A) as well as concentrations of Na^+^ (B) and K^+^ (C) measured in the secreted droplets, rates of Na^+^ (D) and K^+^ (E) ion flux expressed independently of water flux, and the ratio of Na^+^:K^+^ (F) in the secreted fluid. In all cases values presented are mean ± sem. Bars that share a letter within a panel do not significantly differ.

### CAPA peptide quantification in the central nervous system of *D. melanogaster*

We developed a sensitive ELISA for the quantification of *D. melanogaster* CAPA peptides with a linear range of 25 fmol to 25 pmol, which spans three orders of magnitude (Figure 5A). To ensure specificity, we tested cross-reactivity of some structurally-related insect peptides, including pyrokinins (*A. aegpyti* CAPA-PK1: AGNSGANSGMWFGPRL-NH_2_; (Predel et al., 2010)), sNPF (*A. aegpyti* sNPF-1_(4-11)_: SPSLRLRF-NH_2_; (Predel et al., 2010)) and FMRFa (*Rhodnius prolixus* FMRFa: GNDNFMRF-NH_2_; (Sedra and Lange, 2014)), but no cross-reactivity was observed when up to 5 pmol of these peptides were tested (data not shown). Using this ELISA, we were unable to quantify any CAPA peptide in haemolymph extracts from pools of 60 adult females (data not shown). Considering the sensitivity of the *D. melanogaster* CAPA peptide ELISA developed herein with reliable detection down to as low as 25 fmol, this result indicates that the amount of circulating CAPA peptide in the fly haemolymph (∼80 nL) is below 1.25 fmol (equating to an effective titre of less than 15 nM). CAPA material in the nervous system of adult female *D. melanogaster* was quantified using the CAPA peptide ELISA finding the average thoracicoabdominal ganglionic extract from a single fly contains approximately 41 fmol of CAPA-like peptides (Figure 5B), which eluted from the C18 Sep-Pak column in the 20% and 30% solvent fractions. Synthetic *D. melanogaster* CAPA2 peptide processed identically using a C18 Sep-Pak cartridge demonstrated a similar elution profile to the CAPA peptide material in the ganglionic extract. Considering the average CAPA peptidergic material determined per fly along with the previously determined adult female *D. melanogaster* haemolymph volume of ∼80 nL (Folk et al., 2001), the maximum achievable haemolymph titre if all CAPA material is simultaneously released from the nervous system would be ∼512 nM.

**Figure 5.**
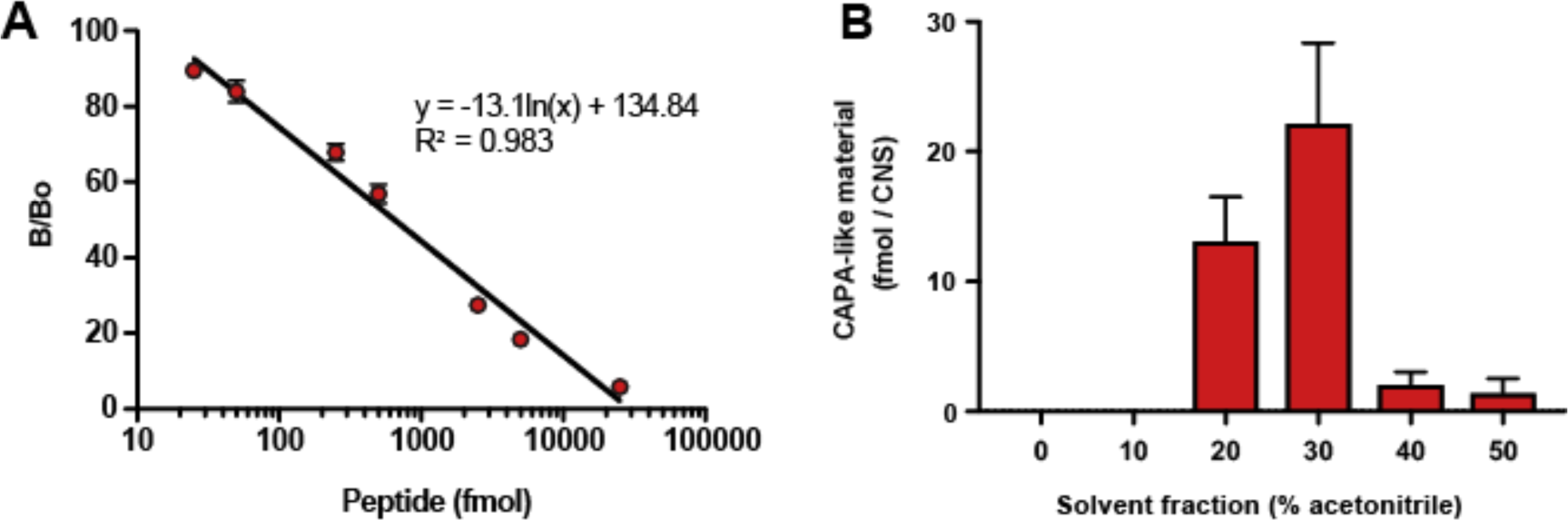
Results of an ELISA against *Drome*CAPA2 in homogenates of the thoracicoabdominal ganglion of female *D. melanogaster*. A) Linear regression analysis of standard curve of ELISA generated using *Drome*CAPA2. Error bars that are not visible are obscured by the symbols. B) Mean ± sem quantity of CAPA-like material found in the thoracicoabdominal ganglion determined from quantification of three biological replicates, each containing 20 ganglia.

## Discussion

This study is the first to report contrasting dose-dependent effects of CAPA peptides on fluid and ion secretion by the Malpighian tubules of *Drosophila melanogaster*, and the first to describe negative effects of a neuropeptide on the chill tolerance of any insect. These results support previous findings related to both high and low doses of CAPA peptides, and raise additional questions about the role of CAPA neuropeptides in insects *in vivo*. Similar to prior evidence (Terhzaz et al., 2015), we found that a micromolar dose of CAPA peptide led to faster recovery from chill coma in *D. melanogaster*. We also report, for the first time, a significant effect of CAPA administration on survival following prolonged chilling (Figure 2B).

Notably, the effects of CAPA injection on chill coma recovery time (CCRT) were similar whether flies were injected early or late in the cold stress (see Figure 1, insert). Recovery from chill coma has been suggested to be dependent on the degree to which an insect has lost ion balance in the cold (dependent on the temperature and duration of cold stress), as well as the rate of ion and water homeostasis recovery following rewarming (MacMillan et al., 2012; Overgaard and MacMillan, 2017). If this is the case, our current result implies that the effects of CAPA peptide on ion and water balance are minimal during the cold stress and that the titre of CAPA in the haemolymph is instead primarily influencing rates of ion transport (and thus the recovery of ion balance) upon rewarming. Given these results, and the knowledge that CAPA receptors are located exclusively in the Malpighian tubules of *D. melanogaster* (Terhzaz et al., 2012), we considered the effects of CAPA peptide on Malpighian tubule function at room temperature that is most relevant to its effects on chill coma recovery. It is possible that the peptide simply cannot bind at low temperatures or otherwise does not alter tubule function in the cold. Alternatively, this result may simply support the observation that rates of transport are strongly suppressed during chilling (approx. 20-fold between 25°C and 0°C in the same population of flies used in the present study (Yerushalmi et al., 2018)), and as such, stimulation or further suppression is unlikely to have any measurable effect. Addressing these possibilities will require careful analysis of the effects of temperature on neuropeptide signaling and renal function across a range of temperatures, as the vast majority of neuropeptide effects in ectotherms have been documented at or near room temperature (De Haes et al., 2015; Nässel and Winther, 2010).

Injection of CAPA peptide was previously demonstrated to have no effect on the survival of *D. melanogaster* following 1 h at −6°C (Terhzaz et al., 2015). Our approach in the present study differed in that the flies were instead subjected to a chronic exposure to a less extreme temperature (16 h at 0°C). Here, flies injected with a low dose of CAPA (10^−15^ M) 30 mins before they were removed from the cold suffered greater chilling injury, while those injected with a high dose (10^−6^ M) were significantly less injured than control flies 24 h following the cold stress. Injuries suffered from chilling in the absence of ice formation are often conceptually divided into direct chilling injury (resulting from severe acute cold stress) and indirect chilling injury (resulting from chronic, but more mild cold stress). These two forms of injury have also been suggested to be associated with different underlying mechanisms; while direct chilling injury is thought to be a consequence of irreversible membrane phase changes and protein denaturation, indirect chilling injury is instead attributed to a more gradual loss of ion and water balance or oxidative stress (Koštál et al., 2006; Overgaard and MacMillan, 2017; Teets and Denlinger, 2013). Our results support the notion that direct and indirect chilling injury are influenced by independent physiological mechanisms, and that neuropeptide effects on ion and water balance may mitigate or exacerbate indirect chilling injury while having little effect on direct chilling injury. Regardless of the mechanisms at play, the effects of CAPA we observed on survival following cold stress mirror our observations for CCRT; while high doses of CAPA improved chill tolerance in *D. melanogaster*, low doses had the opposite effect.

Receptors for CAPA peptides are found only in the Malpighian tubules of *D. melanogaster* (Terhzaz et al., 2012), so we focused our attention on the effects of low and high doses of CAPA on tubule fluid and ion secretion rates using Ramsay assays. Exposure of tubules to femtomolar (10^−15^ M) doses of CAPA peptide reduced rates of ion and fluid secretion by the tubules (Figure 3D,E). Since the initial description of CAPA peptides as modulators of Malpighian tubule secretion rates in *D. melanogaster* (Davies et al., 1995), no studies to our knowledge, have tested the effects of this peptide on fluid secretion below concentrations of 1 nM (10^−9^ M) in this species. Instead, most have described the stimulatory effects of higher concentrations (typically 10^−8^ to 10^−6^ M), and this approach has led to the elucidation of the signaling pathway underlying this effect (Davies et al., 1997; Kean et al., 2002; MacPherson et al., 2001; Pollock et al., 2003; Pollock et al., 2004; Rosay et al., 1997). Our results suggest that CAPA peptide is anti-diuretic at lower (femtomolar to picomolar) concentrations in *D. melanogaster*. Very similar effects of low concentrations of CAPA peptide have been observed in both larval (Ionescu and Donini, 2012) and adult (Sajadi et al., in press) mosquitoes (*Aedes aegypti*). Our results suggest that low concentrations of CAPA impair cold tolerance by slowing rates of ion (particularly K^+^) and water flux through the Malpighian tubules upon rewarming, thereby reducing the speed at which flies can re-establish osmotic and ionic balance following cold exposure.

In contrast to several previous studies on *D. melanogaster* (Davies et al., 1995; Davies et al., 1997; Kean et al., 2002; MacPherson et al., 2001), we found that exposing tubules to micromolar (10^−6^ M) doses of CAPA did not stimulate fluid secretion (Figure 3). We cannot account for this discrepancy. In the present study, however, despite failing to stimulate fluid secretion, 10^−6^ M CAPA instead led to kaliuresis (higher [K^+^] in the secreted fluid). This finding is significant, as it presents a plausible mechanism for increased chill tolerance following injection of high doses of CAPA peptide. In studies conducted on *D. melanogaster* to date, improvements in chill tolerance are associated with increased rates of K^+^ clearance by the tubules. For *D. melanogaster*, acclimation of flies to 10°C is associated with a compensatory increase in the rates of K^+^ secretion by the tubules (Yerushalmi et al., 2018). Similarly, cold-adapted *Drosophila* species maintain high tubule K^+^ secretion rates in the cold (3°C) better than species adapted to warmer climates (MacMillan et al., 2015d). Either of these strategies would help to avoid hyperkalemia in the cold and/or more rapidly recover from ionic imbalance upon rewarming, provided that rates of K^+^ reabsorption are simultaneously kept constant or reduced along the gut epithelia as is the case for both cold acclimated and cold adapted *Drosophila* (Andersen et al., 2017c; Yerushalmi et al., 2018). Thus, although the direct effects of 10^−6^ M CAPA on Malpighian tubule activity observed herein are not in line with previous reports in *Drosophila*, they are internally consistent. We therefore argue that micromolar doses of CAPA peptide improve chill tolerance via kaliuretic activity (i.e. through stimulating K^+^ secretion), with or without stimulating fluid secretion concurrently.

If femtomolar doses of CAPA impair chill tolerance in *Drosophila* and do so by inhibiting fluid and ion secretion in the Malpighian tubules, we predicted that they may do so via the NOS/cGMP/PKG pathway. In larval and adult mosquitoes (*A. aegypti*), low doses of cGMP (10^−9^ to 10^−6^ M) mimic the anti-diuretic effects of low doses of CAPA (10^−15^ M), with maximal inhibition of secretion observed at 10^−8^ M cGMP (Ionescu and Donini, 2012; Sajadi et al., in press). In the case of larval mosquitoes, higher doses of cGMP (10^−3^ M) induce a very modest (non-significant) increase in fluid secretion (Ionescu and Donini, 2012), while in adult mosquitoes no such stimulation could be induced with higher levels of cGMP (Sajadi et al., in press). Accordingly, in the present study we tested whether similar low (10^−8^ M) and high (10^−3^ M) doses of cGMP could mimic these effects in *Drosophila* (Figure 4). While the effects of 10^−8^ M cGMP mirrored the effects of a low dose (10^−15^ M) of CAPA in *Drosophila* (reduced rates of fluid, Na^+^ and K^+^ secretion), 10^−3^ M cGMP did not stimulate fluid secretion, nor did it induce kaliuresis. Indeed, exposure of *D. melanogaster* tubules to 10^−3^ M cGMP had no significant effects on tubule secretion rates, ion concentrations in the secreted fluids, nor rates of ion flux by the tubules (Figure 4). Thus, the impacts of cGMP on tubule secretion in adult *Drosophila* appear to mimic those observed in *A. aegypti* (Ionescu and Donini, 2012; Massaro et al., 2004; Sajadi et al., in press) as well as many other insects including beetles (Eigenheer et al., 2002; Wiehart et al., 2002) and hemipterans (Paluzzi and Orchard, 2006; Quinlan and O’Donnell, 1998). Our results support the idea that low doses of CAPA peptide slow rates of fluid secretion through cGMP signaling, since this second messenger mimicked the anti-diuretic activity of this neuropeptide. In larval *A. aegypti*, stimulatory effects of high doses of AedesCAPA-PVK-1 or high doses of cGMP can be reversed through addition of specific inhibitors of protein kinase A (Ionescu and Donini, 2012), which suggests that high levels of CAPA peptide may be pharmacological, inadvertently activating the signaling cascade that drives diuresis and thereby overwhelming any effects of cGMP. As we did not observe diuretic effects following application of high titres of CAPA peptide in the present study, we were unable to test whether a similar effect can explain CAPA-induced diuresis in *D. melanogaster*.

Given our observations that CAPA peptide can both improve and hinder cold tolerance in *D. melanogaster* depending on the dose applied, we were curious whether flies are capable of reaching micromolar titres of CAPA peptide in the haemolymph. Accordingly, we developed a *D. melanogaster* CAPA peptide-specific ELISA. Despite pooling haemolymph of 60 flies per sample, we were unable to detect CAPA peptides in the haemolymph of *D. melanogaster*, which suggests that total CAPA levels in these samples (collected from flies under control conditions, 23°C) are below our lowest standard (25 fmol). In order for us to detect CAPA in these pooled samples, each fly would have to contribute approximately 0.42 fmol of CAPA, which represents ∼1% of the CAPA peptide quantified in a single thoracicoabodominal ganglion (see below), and our technique of haemolymph extraction typically obtains ∼56 nL of haemolymph from an adult female fly (MacMillan and Hughson, 2014). Thus, in order for us to detect CAPA in the haemolymph, *D. melanogaster* would have to have a mean circulating titre of CAPA peptide equal to or greater than 7.4 nM. As we did not detect CAPA in these samples we suggest that resting titres are below this concentration. Using the same ELISA, however, we were able to detect CAPA peptide in pooled samples of the thoracicoabdominal ganglion (Figure 5), a region of the CNS that houses the Va neurons, where CAPA is produced and stored in *Drosophila* (Kean et al., 2002; Terhzaz et al., 2015). Based on the abundance of CAPA neuropeptides in the entire nervous system where CAPA is produced, we estimate that if all of this peptide was released at once, flies could reach a maximum of about 500 nM CAPA peptide circulating in the haemolymph. This *en masse* release of all CAPA content is unlikely however, since neuropeptides are released as neurohormones from specialized neurohaemal organs (Wegener et al., 2006), including notably the abdominal perivisceral organs where CAPA peptides have been localized and found to be most abundant in a variety of insects (Predel and Wegener, 2006), including in *D. melanogaster* (Predel et al., 2004). Importantly, and consistent with earlier observations in the blowfly *Calliphora erythrocephala* (Duve et al., 1988; Nässel et al., 1988), the *D. melanogaster* adult abdominal neurohaemal organs are directly incorporated into the fused ventral ganglion and localized to the dorsal neural sheath (Predel et al., 2004). In light of this, these results suggest that if flies are capable of reaching micromolar levels of CAPA in the haemolymph, it would likely require, at minimum, a doubling of CAPA peptide abundance above resting levels in the CNS and synchronous release of all CAPA peptides stored within the nervous system. This potential complete release *en masse* is unlikely, however, since *in vitro* induction of neuropeptide release from neurohaemal organs has shown to be only fractional amounts compared to the total immunoreactive material present within the nervous system or neurohaemal organ. For example, in the cockroach *Leucophaea maderae*, leucokinin release from the retrocerebral complex induced by depolarization using high potassium saline resulted in only about 2% release of the total immunoreactive material present within the corpora cardiaca-corpora allata complex (Muren et al., 1993). Similarly, in the house cricket *Acheta domesticus*, release of acheta-kinin following depolarization with high potassium saline from the retrocerebral complex, which is the richest source of this neuropeptide, found released peptide represented less than 4% (i.e. ∼70 fmol released from ∼1800 fmol stored in each retrocerebral complex) of the total acheta-kinin immunoreactive material present within this neurohaemal organ (Chung et al., 1994). Lastly, we note that both cold and desiccation stress have been demonstrated to cause upregulation of Capa mRNA, which may elevate CAPA levels in the CNS, and CAPA has been suggested to only be released in *D. melanogaster* upon removal from the desiccation or cold stress (Terhzaz et al., 2015). Further efforts are thus required to determine whether or not *Drosophila* and other dipterans are capable of reaching levels of CAPA that can stimulate diuresis, and if so, which abiotic conditions specifically lead to this strategy. Critical to this discussion is the direct detection and measurement of circulating levels of CAPA peptide in nanoliter scale haemolymph samples under a variety of highly dynamic conditions, and such an approach in future studies could involve matrix-assisted laser desorption/ionization time-of-flight mass spectrometry (Chen et al., 2009; Fastner et al., 2007).

Chronic exposure to low temperatures suppresses the ability of insects to maintain ion and water homeostasis, causing progressive hyperkalemia and cell death. Our results suggest that CAPA peptides can positively and negatively impact chill tolerance in *D. melanogaster* in a dose-responsive manner. Low (femtomolar) doses of CAPA cause anti-diuresis and limit clearance of [K^+^] at the Malpighian tubules, limiting the ability of flies to recover ion and water balance upon rewarming and impairing chill tolerance. By contrast, high (micromolar) doses of CAPA cause kaliuresis (and based on previous reports also diuresis), facilitating [K^+^] clearance from the haemolymph and improving chill tolerance. We argue that the anti-diuretic effects of CAPA operate through cGMP, and question whether levels of CAPA peptide can reach micromolar levels and stimulate diuresis *in vivo*. Although a wide variety of other neuropeptides are known to influence insect ion and water balance through their effects on Malpighian tubule and gut epithelia, none other than CAPA have been tested in the context of chill tolerance.

## Acknowledgements

The authors wish to thank Carol Bucking for generously providing space in which to conduct some of these experiments.

## Author Contributions

H.A.M., J.P.P., B.N., S.W. G.Y. and L.M. conducted the experiments. H.A.M. and B.N. conducted the data analysis, H.A.M. and J.P.P. drafted the manuscript, and all authors edited the manuscript.

